# Nest traits for the world’s birds

**DOI:** 10.1101/2023.06.06.543860

**Authors:** Catherine Sheard, Sally E. Street, Susan D. Healy, Camille A. Troisi, Andrew D. Clark, Antonia Yovcheva, Alexis Trébaol, Karina Vanadzina, Kevin N. Lala

## Abstract

**Motivation:** A well-constructed nest is a key element of successful reproduction in most species of birds, and nest-building strategies vary widely across the class. Macroecological and macroevolutionary studies tend to group nest design into a small number of discrete categories, often based on taxonomic inference. In reality, however, many species display considerable intraspecific variation in their nest-building behaviour, and broad-level categories may include many functionally distinct nest types. To address this confusion in the literature and facilitate future studies of broad-scale variation in avian parental care, we here introduce a detailed, global comparative database of nest building in birds, together with preliminary correlations between these traits and species-level environmental variables.

**Main types of variable contained:** We present species-level data for nest structure, location, height, material composition, sex of builder, building time, and nest dimensions.

**Spatial location and grain:** Global. Maps are presented at the 1°x1° level.

**Time period and grain:** Included species are generally extant, although we present some data for recently extinct taxa. The data was collected 2017-2021 and was drawn from secondary sources published 1992-2021.

**Major taxa and level of measurement:** Partial or complete trait data is presented for 8,601 species of birds, representing 36 of 36 orders and 239 of 243 families.

**Software format:** Data have been uploaded as Supplementary Material in .xlsx format and are separated by species and source for all traits (S1) as well as summarised at the species level for structure and location variables (S2).

## Introduction

Nest-building behaviour is widespread in birds, and variation in nesting properties is thought to reflect both lineages’ evolutionary histories (Collias, 1997; Fang et al., 2018; Medina et al., 2022; Price & Griffith, 2017) and species’ adaptations to their environments (Collias & Collias, 2014; Deeming & Mainwaring, 2015; Mainwaring et al., 2014). For example, the most speciose order of birds, the Passeriformes (passerines), is believed to have evolved from a cavity-nesting ancestor (Collias, 1997; Fang et al., 2018), with transitions first to dome nests and then to cup-shaped nests (Fang et al., 2018; Price & Griffith, 2017); such an evolutionary history potentially represents trade-offs between the former’s protection from predators and the environment (Hall et al., 2015; Martin et al., 2017; Matysioková & Remeš, 2018) and the latter’s facilitation of niche exploration and modification (Collias, 1997; Fang et al., 2018; Medina et al., 2022; Odling-Smee et al., 2013; Price & Griffith, 2017). Interspecific variation in avian nesting strategy has been linked to a range of ecologically important traits ranging from clutch size (Jetz et al., 2008) and developmental durations (Cooney et al., 2020; Minias & Janiszewski, 2023; Street et al., 2022) to brain structure (Hall et al., 2013) and adherence to Bergmann’s Rule (Mainwaring & Street, 2021), while the relationship between nesting strategy and egg shape (Birkhead et al., 2019; Stoddard et al., 2019; Stoddard et al., 2017) and of nest traits and environmental variation (Englert Duursma et al., 2018; Martin et al., 2017; Medina, 2019; Perez et al., 2020) remains actively debated.

Broad-scale studies of variation in nest morphology and location, however, often aggregate many different nesting behaviours into few, broad categories. For example, Stoddard et al. (2017) scored nest location as ‘non-cavity ground’, ‘non-cavity elevated’, or ‘cavity’ and nest structure as ‘scrape/bed’, ‘plate’, or ‘cup’, neglecting for instance the potential impact of domes. Other sources conflate structure and location: Jetz et al. (2008), for example, coded nest type as ‘open’, ‘half-open’, or ‘closed’, while Cooney et al. (2020) coded nest type as ‘cavity’, ‘closed’, ‘open’, or ‘mixed’. Such groupings can obscure the varying ecological costs and benefits of different strategies. For example, while enclosed nests are thought to be associated with increased protection from predators (Lack, 1948), obligate cavity nesters face much stronger competition for nest sites than do facultative cavity nesters (Martin, 1993a; Martin & Li, 1992). A detailed species-level coding system that carefully distinguishes different nest morphologies and strategies would allow for a researcher to discern between the attributes that are biologically relevant to their own set of hypotheses, rather than relying on a system less suited to their needs.

Furthermore, nest behaviour is often assumed to be invariant at higher taxonomic levels. For example, Jetz et al. (2008) inferred ‘nest type’ within genera, whereas Price & Griffith (2017) and Fang et al. (2018) scored nest shape/structure, location, and exposure/placement at the family level. While this strategy is appropriate for some types of questions, taxonomic inference ignores the tremendous intraspecific and intra-taxon variation in nest behaviour that can be found in the world’s birds (Billerman et al., 2022; Collias & Collias, 2014; Hansell, 2000), as well as the many gaps English-language Western science has in its knowledge of tropical natural history (e.g., Lees et al., 2020).

Here, we present a detailed database of nesting traits (structure, location, height, materials, sex of builder, building time, and size) for the world’s birds. We record intraspecific variation where appropriate, and we note both uncertainty in our coding and where we were unable to find species-level information. We also present a phylogenetically corrected summary of major environmental and morphological correlates of key global variation in nest structure and location, as well as an exploration of geographic biases present in our dataset. We hope that this level of precision and broad taxonomic scope will facilitate future studies of the macroecology and macroevolution of avian parental care as well as direct attention to fruitful directions for future behavioural research.

## Data collection

We targeted text descriptions and photographs published in three sources of information: the Handbook of the Birds of the World Alive (2017-2018), Neotropical Birds Online (2019-2020), and the Birds of North America Online (2019-2021), using the BirdLife International taxonomy. Note that these three sources have subsequently been combined into a single resource, the Birds of the World (Billerman et al., 2022), under a different taxonomy, the eBird/Clements checklist. Coding was done by six researchers (CS, SES, CAT, ADC, AY, and AT). Two sets of researchers (CS, CAT, and ADC; CS and AY) were able to meet regularly to mutually resolve any uncertainties to agreement; most of the data collected by the other two researchers (SES and AT) were checked and if necessary re-coded by a second coder (CS). Two researchers (CS and SES) also each spot-checked an arbitrary set of species. In total, 4072 entries (25.5%) were checked by at least one person other than the original coder.

This data collection process at times generated instances of uncertainty, such as due to vague textual descriptions, unclear photos, or information reported in the secondary source as being suspicious to the author of that source. For example, an entry might note that a species nests in a cavity, but it may not be clear whether that species excavates that cavity or not. We therefore introduced a measure of *uncertainty* in our coding scheme, which allowed the coder to mark for the potential presence of a trait.

We included only breeding nests (i.e., the location of the eggs), rather than any other nest-like construction (e.g., display courts, roosting sites). Our fine-grained classification also targets where possible the bird’s own actions, allowing researchers to distinguish between cavities constructed by primary excavation (a type of nest structure) and secondary cavity nesters (a type of nest location), as well as between facultative and obligate use of different nest strategies. Further information on each of these variables can be found in the Supplementary Materials.

### Nest structure

Inter-specific variation in nest structure is thought to correlate with differences in protection from predators and the environment (Collias, 1997; Englert Duursma et al., 2018; Mainwaring et al., 2014; Martin et al., 2017; Medina, 2019), as well as facilitate or limit the exploration of new ecological niches (Medina et al., 2022; Odling-Smee et al., 2013). We here distinguish among ten major types of constructed nest structures: *none* (birds that lay their eggs directly onto bare substrate or into pre-existing, unmodified cavities), *scrape* (an open, shallow depression created by the bird, with or without a lining), *platform* (a shallow, flat, or saucer-shaped nest with a constructed base and a central depression), *cup* (a constructed nest with walls and a base), *dome* (an enclosed, roofed nest with a small entrance hole), *dome-and-tube* (a multi-chambered dome, such as a dome plus an internal or external entrance hole, including large communal structures), *excavation* (an enclosed cavity created by the species itself), *cavity modifier* (an enclosed cavity formed by a pre-existing cavity subsequently modified by the species itself), and *excavator-with-nest* (a species that fully or partially excavates a cavity and then constructs a structure inside).

We also present a coding system for four rare types of nest structures: *clearing* (a location cleared of debris but with no depression created), *ring* (a location ringed with material with no depression created), *mound* (a strategy whereby eggs are buried in a mound of material, commonly associated with megapodes), and *purse* (a long, pendant pouch, likely providing protection similar to that of a dome but lacking a roof, found in the Icteridae).

Criteria are largely based on Hansell (2000), and full definitions can be found in the Supplementary Material.

### Nest location

Nest location has been found to correlate with clutch size (Jetz et al., 2008) and potentially with egg morphology (e.g., Birkhead et al., 2019, though see Stoddard et al., 2017, Stoddard et al., 2019), as well as to co-evolve with nest structure (Fang et al., 2018; Hall et al., 2015). We here distinguish between seven major categories of nest locations: *artificial structures* (e.g., fences, roofs, nest boxes), *earthen holes, ground*, elevated *rocks, tree holes*, attached *vegetation* (including a subclassification separating out attachments to bushes, trees, and reeds), and fully or partially submerged in *water*. Criteria are largely based on Hansell (2000), and full definitions can be found in the Supplementary Material.

### Nest height

Nest height is often used as a proxy for predation, with higher nests thought to be less accessible to predators (Lima, 2009; Martin, 1993b). We here present values for the minimum and maximum nest height in meters, where available, and noting that ground nests are sometimes slightly elevated by e.g., grass tussocks.

### Nest materials

The materials used to construct a nest can reflect various physical and mechanical properties, including those known or thought to contribute to offspring survival (Bailey et al., 2014; Bailey et al., 2016; Biddle et al., 2018; Breen et al., 2021; Hilton et al., 2004). We here present a compilation of recorded nest materials, which researchers can then search or score for various properties of interest. For one potential categorisation of these materials, see Sheard et al. (2023).

### Sex of builder

Sex-specific contributions to nest building vary by species, as well as with the stage of the nest building process (Mainwaring et al., 2021; Soler et al., 1998). We present what species-level descriptions we were able to obtain, recognising that researchers may be interested in different aspects of sex-specific building process, and therefore not conflating these varying contributions into a single score.

### Building time

The amount of time necessary to build a nest was rarely reported; for what information we could gather, however, we present the minimum, maximum, and average number of days a species has been recorded as spending building a nest, as a potential measure of parental investment (Medina et al., 2022).

### Nest size

We found very little regularity in the reporting of nest size; other researchers, however, may still find these dimensions useful measures of parental investment. For more information on the global correlates of size of passerine cup nests, including comparisons between textual descriptions and museum species as well as an analysis of inter-versus intra-specific variation in size, please see Vanadzina et al. (2023).

### Data patterns

To visualise spatial variation in nest structure and location, we mapped the proportion of six key nest categories (platform, cup, and dome nest structures; ground, vegetation, and artificial nest locations) at the 1°x1° scale using the 2018 BirdLife International breeding and residential range maps (BirdLife, 2018). The distribution of platform nests was relatively uniform, although highest in the Caribbean (Figure 1a). Cup nests were most commonly found in North American and rarely found in Africa (Figure 1c); by contrast, dome nests were generally concentrated in Africa, Australia, and Southeast Asia and rare elsewhere (Figure 1e). Nesting in artificial locations was strongly biased toward the Northern Hemisphere, especially in major deserts (Figure 1b). Ground-nesting strategies showed a remarkable latitudinal gradient, with increased prevalence towards the poles (Figure 1d; cf. Minias & Janiszewski 2023, who demonstrated this in passerines). Finally, nests attached to vegetation were common throughout the world, except at the highest latitudes and on the Tibetan plateau (Figure 1f).

**Fig. 1:**
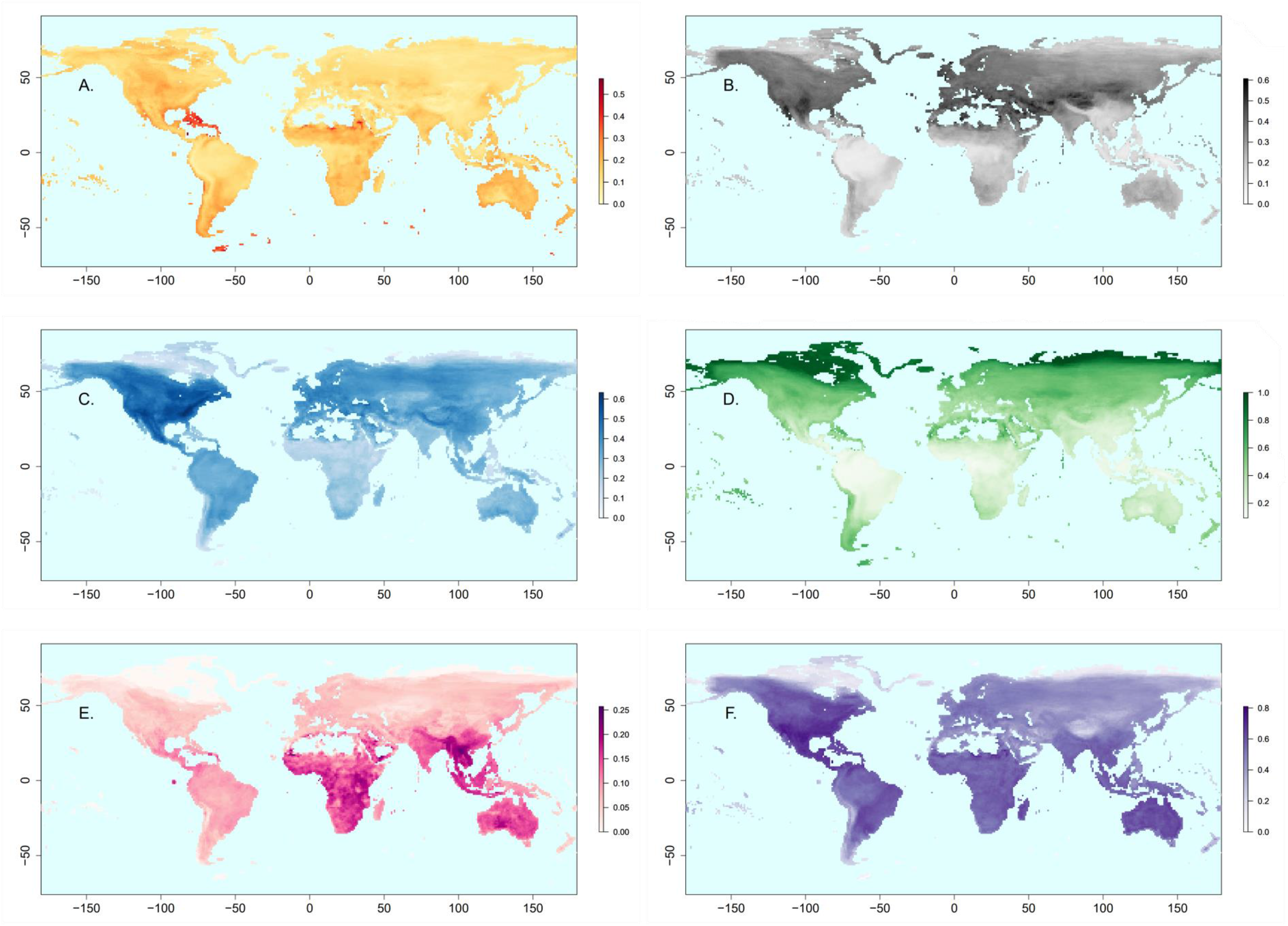
Global distributions of key nest structures and locations. Shown are three nest structures (A. platform, C. cup, and E. dome nests) and three nest locations (B. artificial, D. ground, and F. vegetation nests). Colour shading represents the proportion of each nest structure or location type of species breeding or resident within each 1°x1° square. Squares with < 20 species within our sample (e.g., regions of Antarctica, the Sahara Desert, interior Greenland) have been omitted.

To illustrate exploratory correlations between the most common nest structure and location traits and key environmental variables, we ran Bayesian phylogenetic logistic regressions in the R package *MCMCglmm* (Hadfield, 2010). We first reconciled inter-source variation in nest scores to produce a single species-level set of structure and location scores; for further details on this process, please see the Supplementary Materials. We then obtained species-level values for average breeding range latitude, temperature, precipitation, and annual variability in temperature and precipitation (i.e., temperature and precipitation ‘seasonality’) from Sheard et al. (2020), as well as body mass and a measure of flight ability known as the hand-wing index (HWI; the ratio of Kipp’s distance to the total wing chord) from that source. We included body mass in our models due to the well-established relationship between this variable and avian life history syndromes; we included HWI as it is an increasingly popular proxy for dispersal ability (Claramunt et al., 2012; Kennedy et al., 2016; Pigot et al., 2018; Weeks & Claramunt, 2014) and reflects a key macroevolutionary axis in avian biology, linking for example migratory behaviour, the defence of ecological territories, and diet (Sheard et al., 2020; Weeks et al., 2022).

Models were constructed separately for each of seven nest structure categories (scrape, excavation, platform, cup, dome, dome-and-tube, and none) and seven nest location categories (artificial, earth holes, ground, elevated rocks, tree holes, attached vegetation, and water) as binary response variables and were run across 100 trees randomly chosen from the Hackett backbone of the Jetz et al. (2012) Global Bird Tree. After an initial dummy run to determine start points, each model was run across each tree for a total of 20,000 iterations (burn-in 10,000; sampling rate 1,000; for a posterior sample of 10 per tree). Priors for the fixed effects were set using the command ‘gelman.prior’; priors for the phylogenetic variance were set to *V* = 10^−10^ and *v* = -1, and the residual variance was fixed to 1. To improve output interpretability, all continuous variables were scaled to have a mean of 0 and a variance of 1; body mass and HWI were additionally log-transformed.

Some, though not all, nest structures could be linked with the environment typical of the species breeding range: species were more likely to build cups if they lived in areas with higher precipitation, higher temperature seasonality, and/or lower precipitation seasonality; more likely to build platforms if they lived in areas with higher precipitation seasonality; and less likely to build scrapes if they lived in wetter areas. Furthermore, smaller species were more likely to build domes and more likely to excavate, while larger species were more likely to build platforms or scrapes (Figure 2a). The wing morphology variable HWI was also linked to nest structure; after correcting for mass and environmental correlates, species with high HWI (a proxy for stronger long-distance flight ability) were more likely to build cups, scrapes, or entirely forgo a nest and less likely to build domes or dome-and-tube structures (Figure 2c).

**Fig. 2:**
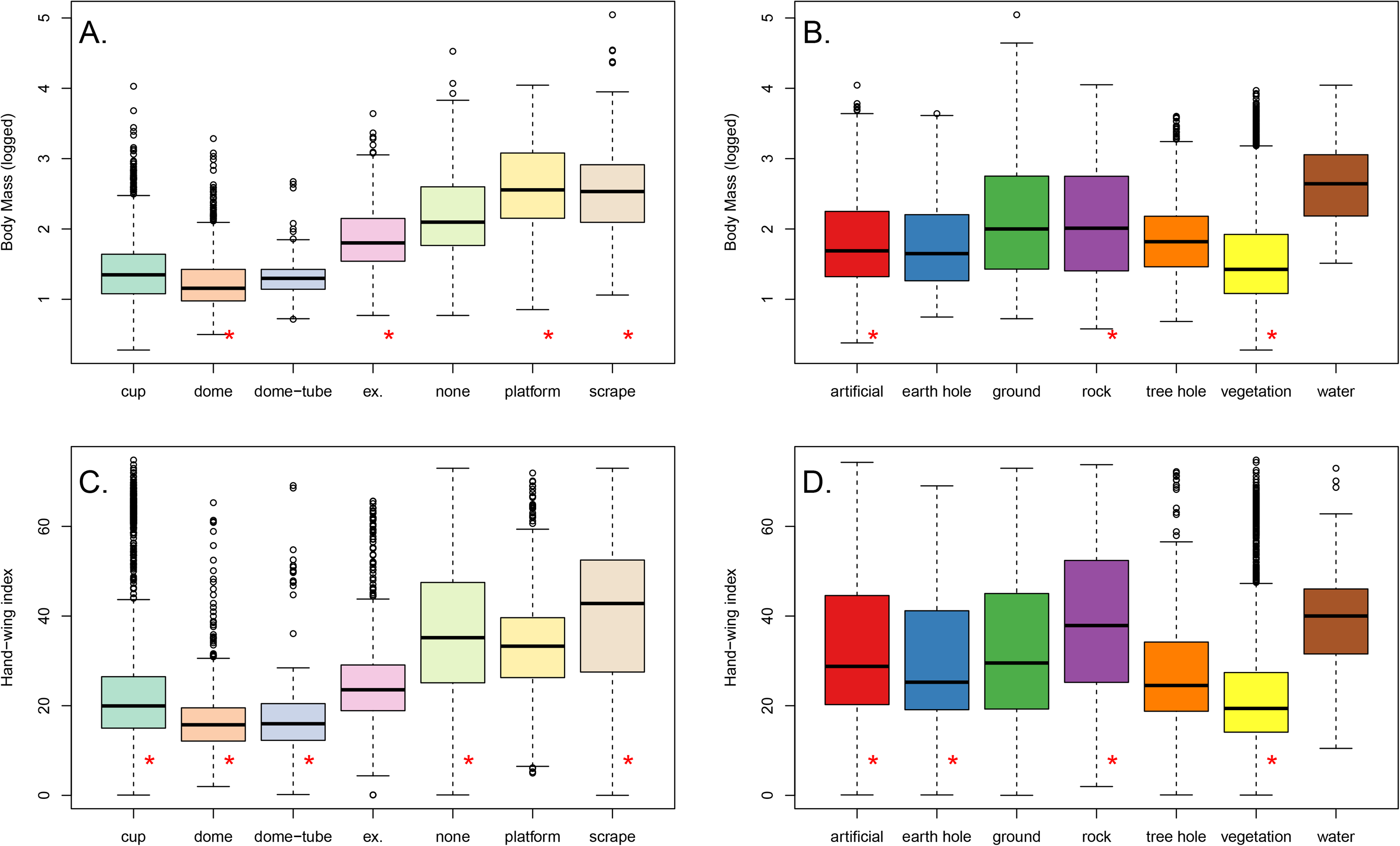
Distribution of body masses (above) and hand-wing indices (HWI, below) for nest structure (left) and nest location (right). Note that some species fall into multiple nest structure/location categories due to intraspecific variation. After correcting for the effects of co-variates and phylogenetic signal, there are significant effects of body mass on the probability that species build dome, excavation, platform, and scrape nests (panel A), as well as nests in artificial, rock, and vegetation locations (panel B), whereas the HWI differences are statistically significant for cup, dome, dome-and-tube, none, and scrape nest structures (panel C), as well as for artificial, earth hole, rock, and vegetation locations (panel D). These statistically significant results are marked with a red asterisk (*). See Tables S1-S7 for full results of the nest structure phylogenetic logistic regressions (panels A, C) and Tables S8-S14 for the nest location phylogenetic logistic regressions (panels B, D). The abbreviations “ex.” = excavations and “dome-tube” = dome-and-tube nests.

Nest location was generally more closely tied to environmental variation than was nest structure. Species were more likely to nest in artificial locations in warmer and/or drier places with greater temperature seasonality and less precipitation seasonality. Species were more likely to nest in earth holes in cooler and/or drier places, and species were more likely to nest in tree holes at higher latitudes, in warmer places, in wetter places, and/or in places with greater temperature seasonality. Species were more likely to nest on the ground in cooler locations, and species were more likely to nest in or near water at lower latitudes and/or in places with greater temperature seasonality. Species were more likely to nest on elevated rocks in cooler, drier, and less seasonal (both temperature and precipitation) places. Species were more likely to nest in vegetation in warmer and/or rainier places with greater temperature seasonality and/or less precipitation seasonality.

Additionally, both heavier species and species with greater HWI (higher flight ability) were more likely to nest in artificial locations or on elevated rocks, while both lighter species and species with smaller HWI (lower flight ability) were more likely to nest in vegetation (Figure 2, panels b and d).

Finally, to explore potential research biases in the nest dataset by geography, we compared the proportion of species lacking nest information across biogeographical realms. We anticipated that species from tropical regions would be most likely to be under-represented in the nest dataset, due in part to pervasive inequalities in the global distribution of research funding (e.g., Lees et al., 2020). Further details on realm scoring and data analysis can be found in the Supplementary Materials. As anticipated, we found that biogeographical realms containing tropical regions generally have more species with missing nest information compared with polar and temperate regions, with species from Oceanian and Neotropical regions particularly under-represented (Figures S1-S4). This is especially notable in the case of builder identity, where the Nearctic realm (i.e., most of North America, plus Greenland) is the only region where >50% of species have documented data.

## Conclusions

We here present a species-level dataset of key nest-building traits for a large sample of birds. Our coding system improves on previous attempts with its level of detail, ability to describe intraspecific variation, and lack of taxonomic inference. We have also described basic environmental and morphological correlations between major structure and location categories, demonstrating that the placement of the nest is more closely linked to broad-scale environmental variation than is the structure of the nest itself. One possible interpretation of this is that environmental factors may have driven finer scaled variation in nest features than gross morphological type (Medina, 2019; Ocampo et al., 2023), such as nest dimensions (Vanadzina et al., 2023); another explanation may be that nest structure is more closely linked to ecological and life history factors not considered here, such as clutch size (Heenan & Seymour, 2011) or predation rates (Collias & Collias, 2014; Hall et al., 2015; Mainwaring et al., 2015; Martin, 1993b; Matysioková & Remeš, 2022).

There are many species about whose nesting strategies Western, English-speaking science knows nothing, particularly in the tropics (Hortal et al., 2015; Lees et al., 2020) (see also Figures S1-S4). We hope, however, that by documenting the variability in nests among well-studied species, we not only provide a useful dataset for future macroecological and macroevolutionary work, but also motivate future detailed documentation of the reproductive biology and behaviour of all the world’s birds.

## Author contributions

SES, CS, KNL, SDH, KV, and ADC designed the data collection protocols. CS, SES, CAT, ADC, AY, and AT collected the initial data. CS and SES checked, cleaned, and analysed the data. CS wrote the manuscript, and SES, KV, SDH, and KNL provided additional writing and editing. All authors approved the final version of the manuscript.

## Acknowledgements

We thank Mike Hansell, Mike Benton, and members of the Healy and Lala labs, especially Sophie Edwards and Helen Spence-Jones, for comments on project design. This work was funded by the John Templeton Foundation (#60501 to KNL) and the European Research Council (788203 ‘Innovation’).

